# Influence of plant growth regulators and nitrogen sources on the production and development of *Brachiaria decumbens* cv. Basilisk

**DOI:** 10.1101/329565

**Authors:** Leone Campos Rocha, Fábio Andrade Teixeira, Márcio dos Santos Pedreira, Daniela Deitos Fries, Evely Giovanna Leite Costa, Ana Cláudia Maia Soares, Daniel Lucas Santos Dias, Abdias José de Figueiredo, Angel Amaral Seixas, Camile Carvalho Pacheco

**Affiliations:** Animal Science Postgraduate Studies Program, São Paulo State University (UNESP), School of Veterinary Medicine and Animal Science, Botucatu, São Paulo, Brazil; Department of Rural and Animal Technology, Bahia State University Southwest (UESB), Itapetinga, Bahia, Brazil; Department of Exact and Natural Sciences, Bahia State University Southwest (UESB), Itapetinga, Bahia, Brazil; Animal Science Postgraduate Studies Program, Bahia State University Southwest (UESB), Itapetinga, Bahia, Brazil; National Postdoctoral Program, Bahia State University Southwest (UESB), Itapetinga, Bahia, Brazil; Animal Science Postgraduate Studies Program, State University of Santa Cruz (UESC), Ilhéus, Bahia, Brazil

## Abstract

The aim was to evaluate the effect of seed treatment with plant growth regulators and nitrogen fertilization methods in *Brachiaria decumbens* cv. Basilisk on the growth and development through germinative characteristics, dry matter yield and physiological composition. Plant growth regulators increased germination and mass yields of plant structures in coated seeds. From the regression estimates, higher germination percentages and root dry mass production were obtained with the use of Plant growth regulators for the initial growth (10.3; 12.8 mL.kg^−1^ seed, respectively). Leaf and pseudostem mass production had a significant interaction effect between seed type and the use of growth regulator. Coated seeds had greater performance with absence or at lower levels of growth regulators, and embryo quality contributed to the greater formation of plant tissues. More efficient levels (8.85 and 9.57 mL.kg^−1^ seed) were observed for the yields of plant structures (leaf and pseudostem). Soil N-fertilization resulted in higher dry mass productions of leaf, stem, shoot and root, as well as for root volume. Rates of photoassimilate were increased by soil fertilization and use of plant growth regulators. The use of exogenous hormonal compounds acts on the organogenesis of plant tissues and increases the development of *Brachiaria decumbens* cv. Basilisk. Soil N-fertilization increase mass yields as it maximizes photosynthetic processes and growth rates.

## Introduction

The shortening of the soil preparation and planting window promoted by the adverse climatic factors reduces the time for recovering degraded pastures. Consequently, greater seed efficiency as regards germination and speed of emergence of seedlings are necessary. Management practices are essential to improve herbage production process using the climatic conditions that favor the development and use of pasture.

The application of synthetic compounds as growth promoters in plants is based on the effect similar to that of phytohormones. Phytohormones (auxins, gibberellins and cytokinins) and the combination of these compounds regulate homeostasis [1]. The PGR acts directly in the signaling network that, through intermediary metabolites, activate genes and then plant responses, such as photosynthesis and the degradation of monosaccharide and proteins.

Growth regulators can potentiate the performance of plants in stress-prone environments, acting on biochemical reactions linked to plant growth. Applications of PGRs based on gibberellic compounds at high levels of active principle per area are shown to be an uneconomical pasture management tool, with reductions in forage mass as application increases. Low concentrations of gibberellin (GA) of 5-10 g.ha^−1^ are enough to positively affect forage yield [2].

PGRs have demonstrated potential applications under stress conditions as they act on the positive regulation of genes linked to the growth and development of plants, increase mechanical impedance in dry soils and improve water use [3]. The use of growth regulators can be applied as an effective technique in plant production, capable of modulating plant responses. However, careful assessment of the synthesis and regulation of these compounds is needed, requiring independent and cross-evaluations among phytohormones’ action.

The use of nitrogen expands plant lifecycle, leading to higher growth rates in the vegetative phase. The increase in photoassimilate production maximizes the structural characteristics of the plant, and allows greater post-defoliation recovery, period in which the plants use energy reserves to reestablish structural and photosynthetic characteristics[4]. Nitrogen is an indispensable constituent in several plant-based organic compounds (amino acids, proteins, nucleic acids) as it increases rates of cell division and expansion [5] and participates in the formation of protoplasm and new cells, as well as stimulating cell elongation [6]. Nitrogen applications to recover degraded pasture areas increase water use efficiency and stimulate plant growth, increasing soil cover [7]. The aim was to evaluate the effect of seed treatment with plant growth regulators (PGR) and nitrogen fertilization methods in *Brachiaria decumbens* cv. Basilisk on the growth and development through germinative characteristics, dry matter yield and physiological composition.

## Material and methods

### Experimental site

The experiments were performed at the State University of Southwestern Bahia (UESB), Itapetinga – BA, between June and September 2017, at the Forage and Pasture Research Center, greenhouse and Laboratory of Anatomy and Ecological Plant Physiology - LAFIEP.

Germination characteristics and initial growth at 30 days after sowing (DAS) were carried out under hermetic storage using a Mangelsdorf germination chamber (MA-401) and seedling trays, respectively. The germination was carried out under a 12-h photoperiod and at constant temperature (28ºC). Growth and development evaluations were performed in a greenhouse, and each pot was filled with eight liters or 12 kg of dry soil, this respective soil volume was used for estimate the area productivity (kg DM.ha^−1^). The soil used was collected at 0-20 cm depth, disintegrated and grounded to pass through a 4-mm sieve.

Analyzes were performed at the Department of Agricultural Engineering and Soils of UESB, and the fertilization recommendations adopted were based on the methodology of the Soil Fertility Commission for the State of Minas Gerais, 5th Approximation [8]. The average, minimum and maximum temperatures and relative humidity within the greenhouse were recorded using a digital thermo-hygrometer, as described in Fig 1.

**Fig 1.**
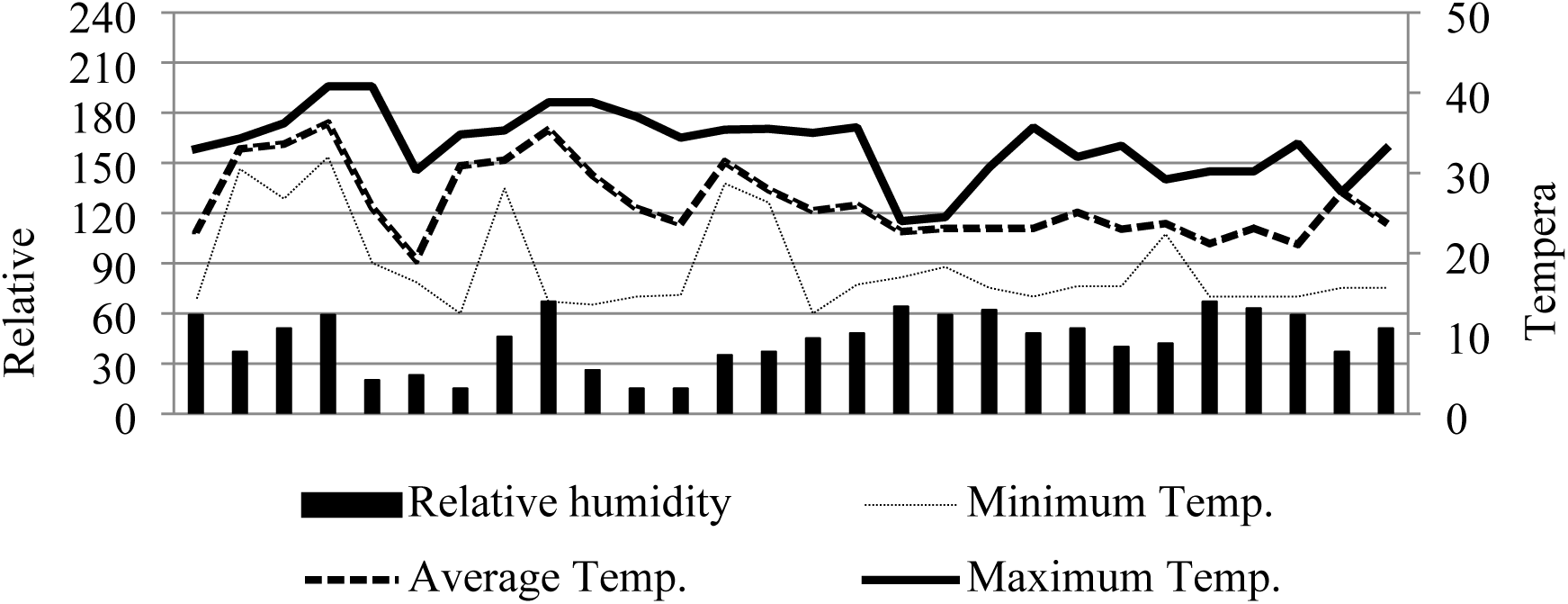
Relative humidity (%) and average, minimum and maximum temperatures (°C) during the experimental period.

### Experimental design

A 2 × 4 factorial scheme was used in a completely randomized design (CRD). Two commercial seed types (coated and conventional) and four levels of the growth regulator (0, 8, 16, 24 mL.kg^−1^ of seed) were used according to the manufacturer’s recommendation.

After the imbibition of *Brachiaria decumbens* cv. Basilisk seeds at different levels of PGR for approximately two hours, the seeds were wrapped in filter paper moistened with distilled water, using 25 seeds per replicate and four replicates per treatment. The PGR was composed of auxin (0.05 mg.L^−1^), cytokinin (0.09 mg.L^−1^) and gibberellin (0.05 mg.L^−1^). For the initial growth assessments (30 DAS) after seed treatment, sowing was performed in a seedling tray with 32 cells per replicate and two replicates per treatment, using as a substrate the sterilized coconut residue to maintain constant humidity throughout the experimental period.

A 2 × 4 factorial scheme was used in a completely randomized design (CRD). The effect of untreated and treated seeds with PGR (0 mL.kg^−1^; 24 mL.kg^−1^) and four nitrogen fertilization methods (no fertilization, soil fertilization, foliar fertilization and multiple fertilization from the combination of soil + foliar fertilization) were evaluated. Eight treatments were used, four plants per replicate and four replicates per treatment, totaling 32 experimental units.

At 30 DAS, a standardization cut was performed adopting an average residue height of 15 cm. At this stage, data collection started, followed by two periods of 28 days. Fertilization was performed after the standardization cut, and the recommendations used were 75 kg.ha^−1^ N and 3 L.ha^−1^ for soil and foliar nitrogen fertilization, respectively. The composition of the foliar fertilizer is described in Table 1.

**Table 1.** Composition of foliar spray

| Density (g.mL<sup>-1</sup>) | Levels (%) |  |  |  |  |  |  |  |  |  |
| --- | --- | --- | --- | --- | --- | --- | --- | --- | --- | --- |
|  | N | P<sub>2</sub>O<sub>5</sub> | Mg | S | Zn | Mn | Cu | B | Mo | Co |
| 1,37 | 2,5 | 15,0 | 1,0 | 1,5 | 3,0 | 2,0 | 0,3 | 0,2 | 0,1 | 0,1 |

Source: Manufacturer.

## Statistical Analyses

Data were analyzed using the MIXED procedure of SAS (SAS Inst. Inc., Cary, NC) with significance declared when *P* ≤ 0.05. Seed, PGR and interaction between effects were analyzed as a fixed effect for all measurements in Exp. 1. Single-degree-of-freedom contrast statements were constructed to analyze PGR effects: linear and quadratic effect of PGR levels. The maximum point was estimated from a regression equation. PGR, Fertilization and interaction between effects were analyzed as a fixed effect for all measurements in Exp. 2. Tukey test was used as a comparison of means with significance declared when *P* ≤ 0.05.

## Evaluations

### Germination

Daily counts of germinated seeds were carried out for 14 days to determine germination percentage and germination speed index (GSI) [9].

### Dry mass production

At the end of the experimental period, the whole-plant yield was determined, followed by desiccation (root, stem and leaf) to determine the fresh mass (DM) and dry mass (DM) of these structures according to [10].

### Chlorophylls and carotenoids

At the end of each experimental period, the third fully expanded leaf of each repetition was collected at 10 a.m. Leaves were cut and an aliquot of 200 mg of fresh mass was immediately prepared [11] by adding 5 mL of dimethyl sulfoxide (DMSO) wrapped in aluminum foil.

After 72 hours, readings at wavelengths of 665, 649 and 480 nm using absorption spectrophotometry were done to quantify the pigments by [12]. The values were adjusted to mg.g^−1^ of fresh mass as follows:

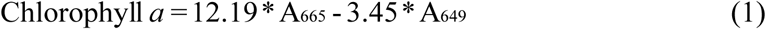

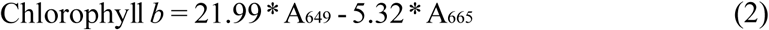

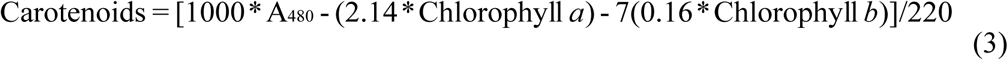

Where A = absorbance. Based on these data, chlorophyll *a*/*b* ratio and total chlorophyll (chlorophyll *a* + chlorophyll *b*).

## Results

The interaction between seed type and growth regulator levels was not significant for germination and germination speed index (GSI) of *Brachiaria decumbens* cv. Basilisk seeds. There was a quadratic effect of growth regulator levels on germination regardless of seed type. According to the equation estimated by regression, the treatment of seeds with 10.3 mL.kg^−1^ promotes greater germination. There was a difference for seed type for both characteristics, in which coated seeds had higher values (Table 2).

**Table 2.** Germination (%) and GSI of different commercial types of *Brachiaria decumbens* cv. Basilisk treated with PGR

| Item | Seed |  | PGR (mL.kg<sup>-1</sup>) |  |  |  | SEM<sup>a</sup> | P-Value<sup>b</sup> |  |  |
| --- | --- | --- | --- | --- | --- | --- | --- | --- | --- | --- |
|  | CON | COA | 0 | 8 | 16 | 24 |  | S | R | S x R |
| Germination<sup>c</sup> | 23.0 | 73.8 | 46.6 | 56.0 | 52.5 | 39.0 | 4.9 | <0.0001 | 0.0182 | 0.0744 |
| GSI | 6.3 | 27.4 | 15.7 | 19.7 | 17.6 | 14.5 | 1.9 | <0.0001 | 0.1130 | 0.3551 |

<sup>a</sup>SEM: Standard error mean
<sup>b</sup> $P \leq 0.05$ ; S: Seed error probability; R: Growth regulator error probability; S x R:

For the number of leaves and tillers, and root dry mass production, the interaction between seed type and growth regulator level was not significant. The number of leaves and tillers were influenced by the seed type used, whereas root dry mass production was not affected. Based on the estimated equation, the treatment of *Brachiaria decumbens* cv. Basilisk seeds with 12.8 mL.kg^−1^ of growth regulator resulted in higher root dry mass production (Table 3).

**Table 3.**
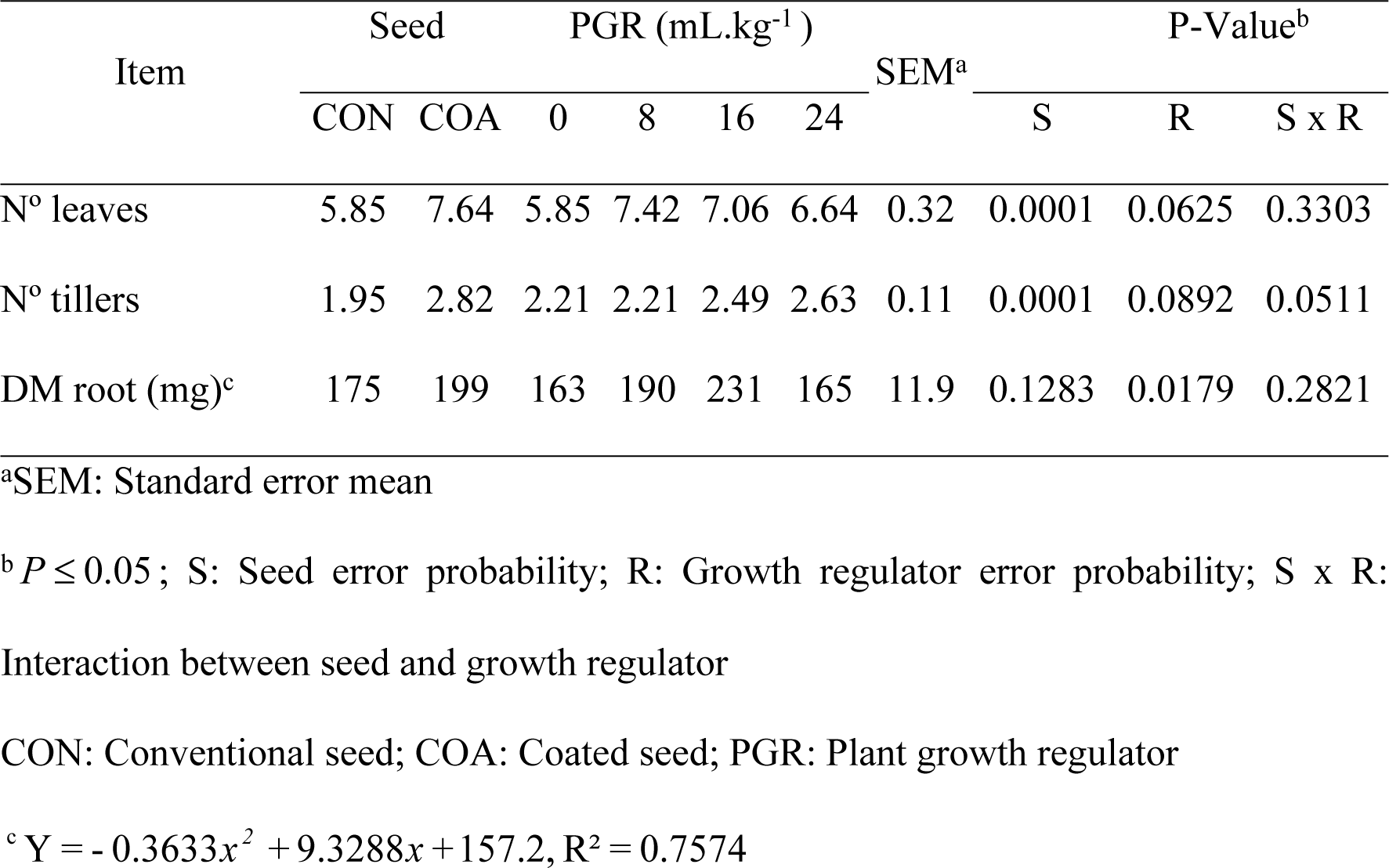
Number of leaves, tillers and root dry mass production in commercial seeds of *Brachiaria decumbens* cv. Basilisk treated with PGR

| Item | Seed |  | PGR (mL.kg<sup>-1</sup>) |  |  |  | SEM<sup>a</sup> | P-Value<sup>b</sup> |  |  |
| --- | --- | --- | --- | --- | --- | --- | --- | --- | --- | --- |
|  | CON | COA | 0 | 8 | 16 | 24 |  | S | R | S x R |
| Nº leaves | 5.85 | 7.64 | 5.85 | 7.42 | 7.06 | 6.64 | 0.32 | 0.0001 | 0.0625 | 0.3303 |
| Nº tillers | 1.95 | 2.82 | 2.21 | 2.21 | 2.49 | 2.63 | 0.11 | 0.0001 | 0.0892 | 0.0511 |
| DM root (mg)<sup>c</sup> | 175 | 199 | 163 | 190 | 231 | 165 | 11.9 | 0.1283 | 0.0179 | 0.2821 |

<sup>a</sup>SEM: Standard error mean
<sup>b</sup> $P \leq 0.05$ ; S: Seed error probability; R: Growth regulator error probability; S x R: Interaction between seed and growth regulator
CON: Conventional seed; COA: Coated seed; PGR: Plant growth regulator

There was an interaction between seed type and the use of growth regulator on leaf and pseudostem dry mass production. In the absence of treatment or at the lowest level tested (8.0 mL.kg^−1^), higher production of leaf and pseudostem dry mass was observed for coated seeds, corresponding to the treatment using 8.85 and 9.57 mL.kg^−1^, respectively. Leaf and pseudostem dry mass production did not differ between seed type at the highest levels tested (16.0 and 24.0 mL.kg^−1^). Conventional seeds required higher levels of the treatment for leaf and pseudostem dry mass production, being 15.5 and 18.1 mL.kg^−1^, respectively (Table 4).

**Table 4.** Dry matter (DM) production of leaf and stem in different commercial seeds of *Brachiaria decumbens* cv. Basilisk treated with PGR

| PGR (mL.kg<sup>-1</sup> ) |  |  |  |  |  |  |  |  |
| --- | --- | --- | --- | --- | --- | --- | --- | --- |
| Seed | 0 | 8 | 16 | 24 | SEM<sup>a</sup> | P-Value<sup>b</sup> |  |  |
|  | DM Leaf (mg) |  |  |  |  | S | R | S x R |
| Conventional<sup>c</sup> | 121b | 184b | 223a | 183a | 14.6 | 0.0001 | 0.0117 | 0.0064 |
| Coated<sup>d</sup> | 254a | 341a | 236a | 223a |  |  |  |  |
| DM Pseudostem (mg) |  |  |  |  |  |  |  |  |
| Conventional<sup>e</sup> | 82b | 123b | 158a | 142a | 12.6 | 0.0001 | 0.0102 | 0.0018 |
| Coated<sup>f</sup> | 176a | 275a | 164a | 163a |  |  |  |  |

<sup>a</sup>SEM: Standard error mean
<sup>b</sup> $P \leq 0.05$ ; S: Seed error probability; R: Growth regulator error probability; S x R: Interaction between seed and growth regulator; PGR: Plant growth regulator
<sup>c</sup> $Y = -0.4009x^2 + 12.449x + 118.41, R^2 = 0.9712$
<sup>d</sup> $Y = -0.3942x^2 + 6.9779x + 268.59, R^2 = 0.5305$
<sup>e</sup> $Y = -0.2184x^2 + 7.9294x + 80.585, R^2 = 0.9665$
<sup>f</sup> $Y = -0.3918x^2 + 7.5044x + 192.48, R^2 = 0.4149$

The interaction between the use of PGR and nitrogen fertilization methods was significant for dry mass production of leaf, stem, shoot, and root. Dry mass production of stem, leaf, and shoot was greater when the use of PGR and soil fertilization were combined, although soil fertilization resulted in higher yields for the evaluated structures regardless of PGR. The use of PGR reduced leaf, stem, and shoot production when combined with multiple fertilization (soil + foliar), differing from the absence of PGR, which was similar to soil fertilization when submitted to multiple fertilization. The use of PGR increased root dry mass production only when combined with multiple fertilization; for the other fertilization types, no influence of growth regulators was observed (Table 5).

**Table 5.**
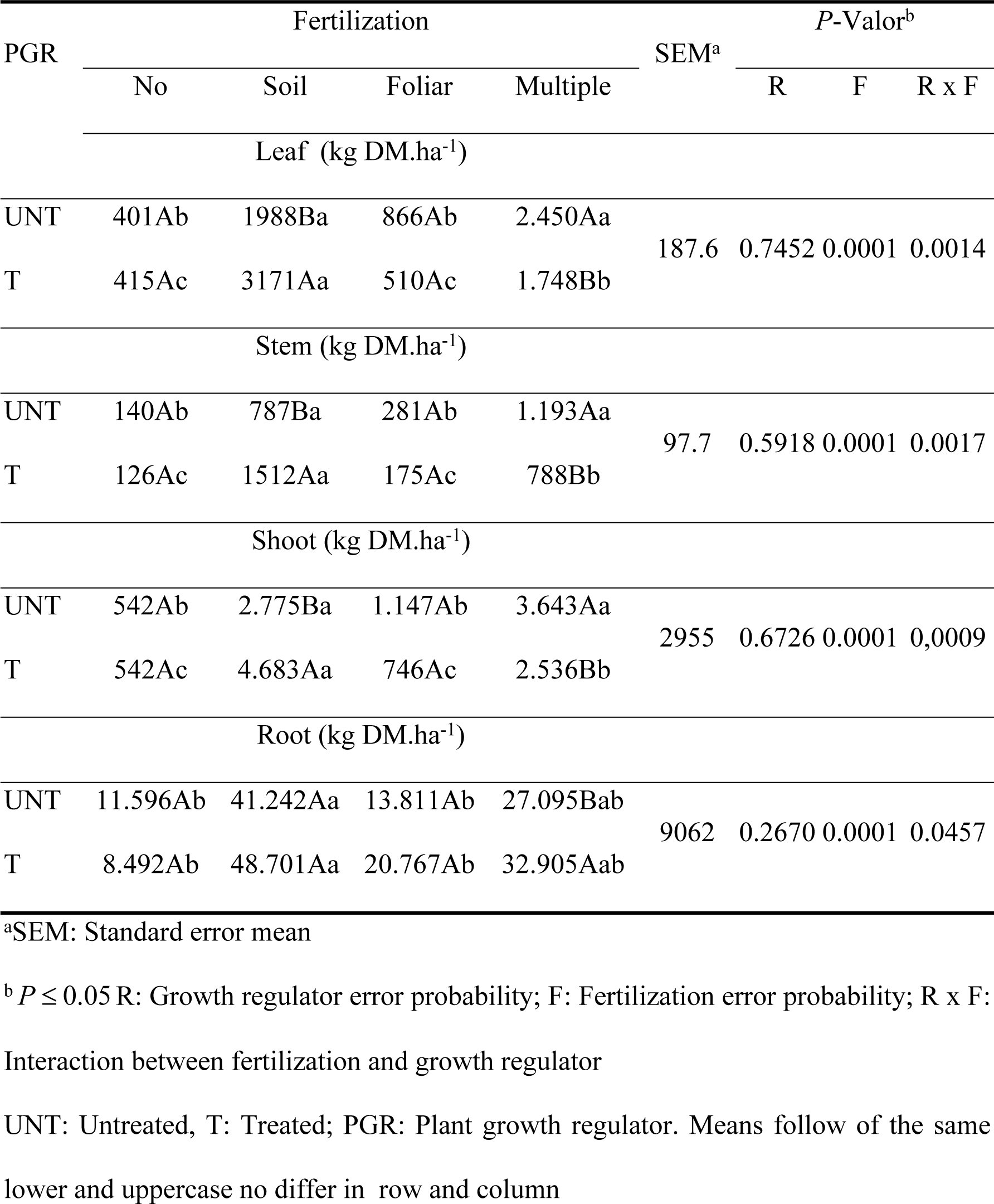
Dry matter (DM) production of leaf, stem, shoot and of *Brachiaria decumbens* cv. Basilisk treated or not with PGR and nitrogen fertilization methods

| PGR | Fertilization |  |  |  | SEM<sup>a</sup> | *P*-Valor<sup>b</sup> |  |  |
| --- | --- | --- | --- | --- | --- | --- | --- | --- |
|  | No | Soil | Foliar | Multiple |  | R | F | R x F |
|  | Leaf (kg DM.ha<sup>-1</sup>) |  |  |  |  |  |  |  |
| UNT | 401Ab | 1988Ba | 866Ab | 2.450Aa | 187.6 | 0.7452 | 0.0001 | 0.0014 |
| T | 415Ac | 3171Aa | 510Ac | 1.748Bb |  |  |  |  |
| Stem (kg DM.ha<sup>-1</sup>) |  |  |  |  |  |  |  |  |
| UNT | 140Ab | 787Ba | 281Ab | 1.193Aa | 97.7 | 0.5918 | 0.0001 | 0.0017 |
| T | 126Ac | 1512Aa | 175Ac | 788Bb |  |  |  |  |
| Shoot (kg DM.ha<sup>-1</sup>) |  |  |  |  |  |  |  |  |
| UNT | 542Ab | 2.775Ba | 1.147Ab | 3.643Aa | 2955 | 0.6726 | 0.0001 | 0,0009 |
| T | 542Ac | 4.683Aa | 746Ac | 2.536Bb |  |  |  |  |
| Root (kg DM.ha<sup>-1</sup>) |  |  |  |  |  |  |  |  |
| UNT | 11.596Ab | 41.242Aa | 13.811Ab | 27.095Bab | 9062 | 0.2670 | 0.0001 | 0.0457 |
| T | 8.492Ab | 48.701Aa | 20.767Ab | 32.905Aab |  |  |  |  |

<sup>a</sup>SEM: Standard error mean
<sup>b</sup> $P \leq 0.05$ R: Growth regulator error probability; F: Fertilization error probability; R x F: Interaction between fertilization and growth regulator UNT: Untreated, T: Treated; PGR: Plant growth regulator. Means follow of the same lower and uppcase no differ in row and column

**Table 6.** Content of chlorophyll *a*, chlorophyll *b*, total chlorophyll, carotenoids and chlorophyll a / b ratio of *Brachiaria decumbens* cv. Basilisk treated or not with PGR and nitrogen fertilization methods

| Item | PGR |  | Fertilization |  |  |  | SEM<sup>a</sup> | P-Valor<sup>b</sup> |  |  |
| --- | --- | --- | --- | --- | --- | --- | --- | --- | --- | --- |
|  | UNT | T | No | Soil | Foliar | Multiple |  | E | A | E x A |
| Chlorophyll *a* | 0.91 | 1.01 | 0.86b | 0.93b | 0.89b | 1.15a | 0.03 | 0.0340 | 0.0002 | 0.5154 |
| Chlorophyll *b* | 0.25 | 0.23 | 0.21b | 0.22ab | 0.24ab | 0.30a | 0.01 | 0.5768 | 0.0191 | 0.2566 |
| Total chlorophyll | 1.16 | 1.25 | 1.06b | 1.15b | 1.14b | 1.46a | 0.03 | 0.1570 | 0.0001 | 0.4016 |
| Carotenoids | 0.44 | 0.47 | 0.44a | 0.44a | 0.46a | 0.48a | 0.007 | 0.0581 | 0.0939 | 0.0939 |
| *a/b* ratio | 3.84 | 4.31 | 4.08a | 4.17a | 4.12a | 3.93a | 0.14 | 0.1110 | 0.9354 | 0.2665 |

<sup>a</sup>SEM: Standard error mean
<sup>b</sup> $P \leq 0.05$ ; R: Growth regulator error probability; F: Fertilization error probability; R x F: Interaction between fertilization and growth regulator
UNT: Untreated, T: Treated; PGR: Plant growth regulator. Means follow the same letters no differ

## Discussion

The use of exogenous hormone compounds changes hormonal balance and biosynthesis levels of their respective metabolites, with a significant increase in auxin, cytokinin and gibberellin levels after 24 h of imbibition [13]. The use of growth regulator increased germination percentages for both seed types, contrasting studies where only cytokinin and auxin sets of formulation were evaluated, with no significant responses [14,15].

The increased concentration of exogenous gibberellin implies greater intracellular transport of intermediary metabolites until the formation of bioactive gibberellin [16,17], which acts on genes responsible for expressing germinative characteristics, increasing the growth potential of embryo and overcoming the mechanical barrier by weakening structures that cover the radicle [15]. The application of a combined hormone regulator brings cross-responses between dormancy and germination when there is a modification of gibberellin/abscisic acid relationships. The use of gibberellic acid (gibberellin) increases germination by reducing the concentrations of abscisic acid, the main promoter of seed dormancy [18]. Abscisic acid regulates the mobilization of storage lipids in the endosperm. Therefore, by decreasing this content, lipid degradation occurs in cotyledons [19].

The absence of gibberellin elicits responses similar to the use of the respective antagonistic hormones, requiring an evaluation beyond the absolute values due to the interaction between hormones [14].

Coated seeds had higher germination percentages and GSI due to their quality. Seed coating provides protection against mechanical damage and nutrients to the embryo [20], favoring greater radicle emergence and seedling emergence rate.

The use of growth regulator did not result in higher number of leaves and tillers, as reported by [21]. Higher root dry mass production was observed because the use of auxin in the composition of the growth regulator increases rhizogenesis. Auxin stimulates stem cell multiplication due to the accumulation of auxin in quiescent centers (QC), increasing the number of root cells [22]. Auxin transport occurs in small quantities by the phloem, but the regulation of the processes involving the auxin is based on the basipetal polar transport, i.e., in the protonated (IAAH) or anionic form (IAA-).

The PIN proteins are bound to plasma membranes enabling the intracellular transport of auxins [17,22,23]. They are located in the endoplasmic reticulum and other endomembranes, contributing to the homeostasis of auxins until the signal response [24]. Evaluating different auxins (IBA, IAA, NAA and BAP) found higher rooting ability (72.4% and 9.25 for frequency of rooting and roots per tiller, respectively) using IBA (Indole 3-butyric acid), the same compound tested [25]. The effectiveness of the use of auxin compounds (IAA, IBA and NAA) in rooting is also proposed by [26], reporting that the use of these compounds resulted in greater rooting percentage, reinforcing the effect on the rhizogenesis that auxin exerts in zones of growth and cellular elongation that affect root growth [27]. The interaction auxin/cytokine regulates processes of formation and maintenance of meristems [28], and both may act separately in the expression of organ-forming genes or interact in an antagonistic way during embryogenesis [29]. Auxins and cytokinins are the main hormones involved in signaling the formation of the shoot apical meristem (SAM), indicating the importance of these proportions and the potential cross-response between these two hormones in the formation of plant organs.

Conventional seeds required greater use of growth regulators because of their reduced efficiency in its use. The best responses observed in the combination of coated seeds and lower levels are justified because they are pure seeds, possibly with higher embryo quality. The coating favored higher yields of growth structures (root and emergence) as mentioned above. When treated with low levels, conventional seeds had lower efficiency of hormone use, suggesting an evaluation of Pure Live Seeds (PLS) that certainly improves their use efficiency.

The evaluations of dry mass production should be analyzed beyond absolute values of the phytohormones used, such as the hormonal interaction, which is comprised of a signaling network, as stated by [27]. Hormones play key roles in promoting growth or stress responses, and each hormone acts on physiological processes and plant development, in which every process is co-regulated by multiple hormones.

The interaction between auxin and cytokinin positively influences plant growth and development by acting on meristems, in the formation of leaves and roots [30]. Gibberellins produced in young leaves in expansion play a primary role in stimulating the auxin reaction, directly affecting the internode elongation [31]. This cross-effect is due to the participation of intermediate receptors of one hormone interacting with components of another.

Higher dry mass yields were observed for all evaluated structures (leaf, stem, shoot, and root) receiving soil fertilization, justifying nitrogen relationship and its greater efficiency to influence photosynthetic processes when absorbed by root.

Greater yields were observed when PGR and soil fertilization were combined, supporting the idea that greater nitrogen supply combined with greater cellular acceleration and synthesis of hormonal metabolites provided by PGR reflects in higher dry mass accumulation (DM) of the evaluated structures. The increased leaf DM production consolidates the improvement of photosynthetic processes that enhanced vegetal response, where the use of PGR resulted in higher levels of chlorophyll *a*, responsible for the photochemical phase of photosynthesis, promoting the formation of ATP in the photophosphorylation process. Corroborating with [32], the use of nitrogen fertilization (50 kg N.ha^−1^) combined with sprays of gibberellin increased DM yields. It is worth emphasizing the influence of the hormonal compound used in the formation of structures such as leaf, stem, and root, already mentioned in experiment I.

The increase in the contents of chlorophyll *a* with the use of growth regulator, combined with the result of mass yield, justifies the efficiency of use of exogenous compounds in the regulation of plant development as they exert functions in the signaling network of photosynthetic processes. The higher production of chlorophyll *a,* a pigment that makes light perception possible in great extent, favors the synthesis of ATP in chloroplasts, being used in the process of photophosphorylation [17]. Consequently, higher energy content is captured by the plant and, therefore, more substrate will be available for plant development.

Little is known about the role of auxins and cytokinins in the regulation of photosynthetic processes. Auxins are linked to processes of phototropism and movement of chloroplasts in the perception of light. In a still inconclusive study, [33] found that auxin may indirectly influence the movement of chloroplasts, an effect caused by the auxin homeostasis disorder. [17] report that the use of growth regulators in the form of plant herbicides (growth regulators) acts by blocking the photosynthetic electron flux, altering the formation of ATP in chloroplasts.

The use of foliar nitrogen fertilization increased the production of chlorophyll *a*, chlorophyll *b* and total chlorophyll, corroborating with [34], who observed increased concentrations of chlorophyll in leaves of plants that received foliar sprays of urea. The availability of nitrogen supplied through foliar application may participate in cytokinin synthesis, which are important for promoting chlorophyll synthesis and delayed senescence. Foliar fertilization allows faster access to nitrogen by growing plant tissues.

## Conclusion

The use of growth promoters (PGR) increases the formation of plant tissues and acts in the promotion of germination and development of structures, making it a key tool in pasture establishment. Lower levels of PGR were recommended in coated seeds when compared to conventional methods. No influence of foliar fertilization was observed when used exclusively, but soil N-fertilization promotes an increase in mass yield and chlorophyll content.

## Acknowledgments

Authors are grateful to Bahia State University Southwest (UESB).

